# Early-Life Exposure to Warming Enhances Sea Urchin Heatwave Tolerance but Fails under Extreme Stress

**DOI:** 10.64898/2026.05.26.728038

**Authors:** L.C. Bonzi, S. Suresh, J. Sourisse, M. Cutracci, A. Chung, D. Romeo, J. Kang, D. Desantis, B.P. Pereira, E. Otjacques, J.R. Paula, T. Repolho, C. Schunter

**Affiliations:** Swire Institute of Marine Science, School of Biological Sciences, The University of Hong Kong, Hong Kong, Hong Kong SAR, China; Department of Environmental Conservation, University of Massachusetts, Amherst, MA 01003, USA; School of Marine Sciences, State Key Laboratory of Marine Resources Utilization in South China Sea, Hainan University, Haikou, Hainan, P. R. China; MARE – Marine and Environmental Science Centre & ARNET – Aquatic Research Network, Faculdade de Ciências da Universidade de Lisboa, Laboratório Marítimo da Guia, Cascais, Portugal; Departamento de Biologia, Faculdade de Ciências da Universidade de Lisboa, Campo Grande, 1749-016, Lisbon, Portugal

**Keywords:** acclimation, pre-exposure, invertebrate, plasticity, climate change, physiology

## Abstract

The exposure to environmental stressors early in life can shape organisms to express more tolerant phenotypes to the same conditions during adulthood, a process called developmental plasticity. However, this acquired acclimation ability might depend on the intensity of the stimulus perceived later in life. Here, we took advantage of a purple sea urchin *Paracentrotus lividus* population developed at abnormally high sea temperature in the proximity of a power plant to test the limits of their developmentally acquired plasticity to increased water temperature. We simulated two marine heatwaves, a category I (moderate) and IV (extreme), and exposed the power plant population as well as a naive population developed in natural sea conditions. We measured their respiration rate and molecular responses to these two heatwaves. Regardless of the population of origin, sea urchins exposed to heatwaves showed higher oxygen consumption, indicating an increase in metabolic rates. At the molecular level, the biggest difference between the two populations was found following the moderate heatwave. Compared to the developmentally acclimated sea urchins, the naive population expressed genes coding for proteins with stress response, chromatin remodeling and RNA splicing functions, while suppressing immune response, revealing that developmental exposures can aid in priming the responses of adults to moderate temperature increases. However, a stronger heatwave leveled the differences between the two populations, with sea urchins from both locations expressing genes involved in proteostasis and detoxification. Nevertheless, regardless of the simulated marine heatwave intensity, sea urchins from the naive population always showed enrichment of the spliceosome pathway compared to power plant urchins, which activated immune response genes instead, reflecting fundamentally different thermal stress-coping strategies shaped by their developmental environments. Overall, our results demonstrate the critical yet context-dependent role of developmental plasticity in shaping the resilience of marine ectotherms to climate change.

## Introduction

Anthropogenic climate change is driving both a rise in average ocean temperatures (i.e. ocean warming) and an increase in the length, frequency and intensity of extreme thermal events, such as marine heatwaves (Frölicher et al., 2018; MHWs; Oliver et al., 2018). MHWs are defined as prolonged periods of abnormally warm ocean temperatures (Hobday et al., 2016; IPCC, 2021), and their occurrence can pose unprecedented thermal stress on marine organisms, with detrimental effects at the individual, population and ecosystem level (reviewed in Smith et al., 2023), including mass mortality events (Garrabou et al., 2022; Hughes et al., 2018). Because of the intrinsic unpredictability and intense nature of MHWs, organismal and population responses to such extreme events are likely to differ from those to steady warming over multiple decades. Instead of geographical range shifting or genetic adaptation to the new thermal niche, for example, acclimation through phenotypic plasticity will potentially have a larger relevance as a coping mechanism (Harvey et al., 2022), especially in organisms with limited movement ability like many marine invertebrates.

Plastic responses to thermal stress could particularly be effective when organisms have already experienced increased temperature in their lifetime, especially during early-life stages. Early exposure to stressors can shape adult phenotype (i.e., developmental plasticity) and prime individuals to better withstand future stress (West-Eberhard, 2003). In fact, aquatic ectotherms, such as marine invertebrates, show higher developmental plasticity compared to other taxa (Pottier et al., 2022). For instance, in the copepod *Acartia tonsa*, development at higher temperature increases thermal tolerance of adults (Ashlock et al., 2024). Similarly, juvenile *Loxechinus albus* sea urchins exposed to a 5°C temperature rise significantly increased both their critical thermal minimum and maximum (Manríquez et al., 2019). Therefore, exposure to warming conditions at early-life stages might enhance marine organisms’ resilience to future thermal stresses. Nevertheless, the adaptive potential of developmental plasticity in changing thermal environments has been shown to have limitations. For example, a recent meta-analysis found that developmental thermal plasticity can vary with ontogeny and heating rate (Pottier et al., 2022), and exposure duration also affects individual future performances (Spinks et al., 2019). Moreover, the adaptive potential of developmental plasticity likely depends on the magnitude of subsequent stressors experienced later in life, with benefits diminishing as the environmental conditions deviate further from early-life experiences (i.e., mismatch between phenotype and environment), or when physiological thresholds are exceeded. Therefore, determining the extent to which developmentally acquired plasticity can buffer against acute or extreme stressors later in life is key for predicting species resilience to thermal stress imposed by MHWs.

Sea urchins are ecologically important grazers that shape ecosystem dynamics and function (Uthicke et al., 2009), and they also represent a valuable wild fishery resource worldwide (Andrew et al., 2002). They possess a pelagic larval stage that allows them considerable dispersal abilities (Barrier et al., 2024; Duran et al., 2004), and juvenile and adult benthic forms which, on the other hand, have limited movement and thus need to rely on plastic responses to cope with environmental changes. Sea urchin benthic stages have been found to be quite tolerant to increased temperatures, being able to maintain their performance until thermal limits are reached (Lang et al., 2023), at which point metabolic depression and mortality usually occur. Indeed, adult sea urchins exposed to warming appear to suffer limited effects on biological responses such as growth, feeding or movement (Lang et al., 2023), suggesting thermal adjustment at the molecular level. Accordingly, recent studies on species such as the short-spined sea urchin (*Strongylocentrotus intermedius*) and the black sea urchin (*Arbacia lixula*) have revealed transcriptional shifts under warming, including activation of antioxidant defenses, apoptosis, inflammation, immune system, autophagy and endoplasmic reticulum stress pathways (Han et al., 2023; Huang et al., 2026; Liu et al., 2023; Pérez-Portela et al., 2020; Zhan et al., 2019). However, such molecular plasticity is limited by the intensity of warming, with an increase of ≥∼4°C approaching physiological thresholds for many sea urchin species (Byrne & Hernández, 2020). Moreover, increased water temperature can have other cryptic effects, such as detrimental impacts on reproductive capacity. In the purple sea urchin *Paracentrotus lividus*, for example, MHWs damage gonadal tissue (Amato et al., 2025), reducing the organism’s reproductive potential with large-scale detrimental effects on population viability (Yeruham et al., 2020). Finally, the duration of exposure has also been proved to have a significant effect on the resilience to warming, with sea urchins showing greater acclimation abilities with longer exposure times (reviewed in Byrne & Hernández, 2020). Together, these findings highlight the importance of developmental thermal history in shaping resilience to subsequent thermal stress, yet the molecular processes underlying such developmental plasticity and whether any acquired benefits vary with the intensity of future warming events remain poorly understood.

To address this, we focused on the purple sea urchin *Paracentrotus lividus* (Lamarck, 1816), an ecologically and economically important species distributed in the North-Eastern Atlantic Ocean and throughout the Mediterranean Sea. This sea urchin is common in rocky intertidal and shallow subtidal habitats, where the temperature ranges from 10 to 25°C (Boudouresque & Verlaque, 2001), with mass mortality events occurring in parts of its distribution range when 30.5°C water temperature is reached (Yeruham et al., 2015). Here, we leveraged a unique purple sea urchin population inhabiting a coastal area adjacent to a power plant to investigate whether post-metamorphic development under elevated temperature conditions can enhance physiological and molecular resilience to marine heatwaves (MHWs). Importantly, individuals from this population were collected nine months after the shutdown of the power plant, meaning they were no longer experiencing elevated temperatures at the time of sampling. Given their size and developmental history, however, these individuals had developed under chronically warmer conditions, allowing us to specifically assess persistent developmental effects rather than acute thermal acclimation. We explored developmental plasticity limits by exposing individuals from this developmentally acclimated population and a naive population from natural control conditions to two MHWs of different intensity: category I (moderate) or category IV (extreme) and measuring their physiological and molecular responses. Building on previous work linking metabolic plasticity to thermal tolerance (Sokolova, 2013), we hypothesize that developmentally acclimated urchins will exhibit enhanced resilience to moderate MHWs but will eventually converge with the naive population individuals under extreme temperatures due to the exceedance of physiological thresholds. Findings from our study will advance our understanding of the limits of developmental thermal plasticity and aid in assessing the vulnerability of marine invertebrates to current and future ocean warming and associated extreme events.

## Methods

### Animal collection and experimental design

Adult *Paracentrotus lividus* sea urchins (test width > 40 mm; Boudouresque & Verlaque, 2001) were collected by hand and SCUBA diving in September 2021 from two locations on the Portuguese coast: Sines (37°55’33’’N, 8°48’32’’W) and Cabo Raso (38°42’35’’N, 9°29’11’’W; Fig. 1A). Sines region hosted a thermoelectric power plant that operated from the mid 1980s until January 14, 2021 (nine months before sample collection, when the power plant was shut down), resulting in seawater temperatures averaging 2.9°C above nearby coastal areas (Suppl. Table S1). Due to the limited dispersal capacity of *P. lividus* across the surrounding sandy substrate (Boudouresque & Verlaque, 2001), individuals from Sines are presumed to have developed and lived under chronically elevated temperatures throughout their lifespan, until one year before sample collection when the power plant was shut down. Cabo Raso site was selected as control, representing a population of naive sea urchins that developed and lived under natural temperature conditions (Suppl. Table S1).

**Figure 1.**
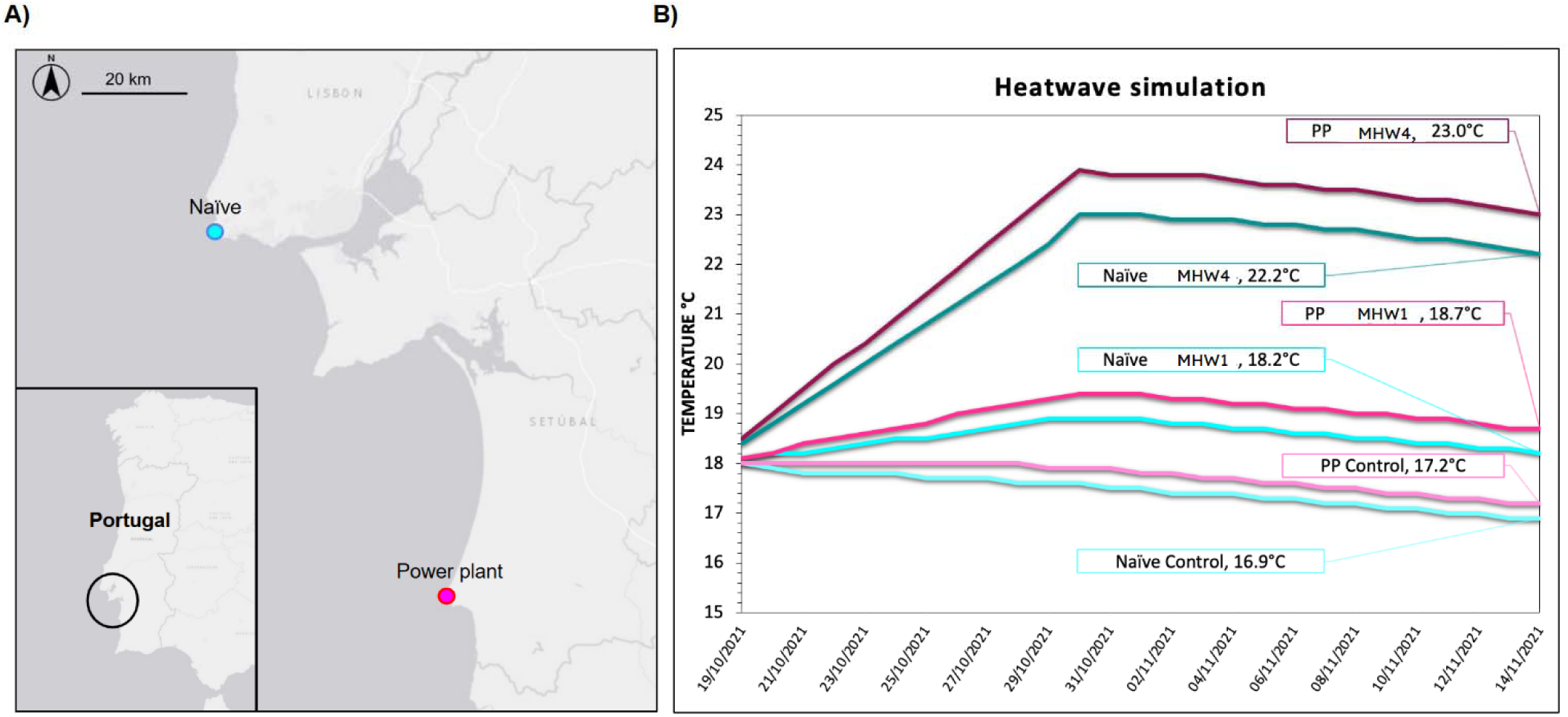
Collection sites and experimental design. A) Map of the Portuguese coast showing the sampling locations of P*aracentrotus lividus*. Blue indicates the naive population collected at Cabo Raso, while pink the power plant population collected at Sines. B) Sea urchins from each population were exposed to control conditions or Marine Heatwave (MHW) simulated category I (“moderate”) and category IV (“extreme”) marine heatwaves. Temperatures shown correspond to the target experimental temperatures reached 14 days after the onset of the heatwave simulations, when sea urchins were subjected to respirometry trials and sampled for molecular analyses. “Naive” refers to the Cabo Raso population and “PP” to the Sines power plant population.

Following collection, sea urchins were transported in aerated seawater to the aquatic facilities of Laboratório Marítimo da Guia (Cascais, Portugal). Here, the animals were weighed, measured (Suppl. Table S2), and acclimated for 27 days in semi-open recirculating aquatic systems supplied with filtered and UV-sterilised natural seawater. Sea urchins were maintained under controlled conditions (17.8 ± 0.1°C; salinity 34.5 ± 0.1 ppt; dissolved oxygen 7.8 ± 0.02 mg L−1−1; pH 8.03 ± 0.01; mean ± SD), fed with *Laminaria ochroleuca ad libitum* and under 14h/10h (light/dark cycle) photoperiod. Seawater parameters were monitored daily, including salinity (HI98319, Hanna Instruments, USA), dissolved oxygen (DO 210, VWR, USA), temperature (DO 210, VWR, USA) and pH (pHenomenal® pH 1100 H, VWR, USA). After acclimation, sea urchins from both populations were assigned to one of three experimental conditions: i) control, where sea urchins were exposed to the water temperature of their respective collection sites; ii) category I marine heatwave (MHWI, “moderate”); and iii) category IV marine heatwave (MHWIV, “extreme”), following Hobday et al. (2018 Fig. 1B; Suppl. Table S3).

Experimental temperatures were determined using 30 years (1991-2021) of sea surface temperature data obtained from NOAA OISST and analysed with R (v4.3.1; R Core Team, 2023) package heatwaveR (v0.0.5; Schlegel & Smit, 2018). Baseline control temperatures were defined according to the local thermal regimes of each sampling location, and marine heatwave (MHW) treatments were simulated relative to these baseline conditions following Hobday et al. (2018). Category I (moderate) and category IV (extreme) heatwaves corresponded to approximately 1–2× and ≥4× the local thermal anomaly above baseline temperatures, respectively. Consequently, experimental temperatures fluctuated throughout the trial following the temporal variation in local SST conditions rather than remaining constant (Fig. 1B). Peak temperatures reached 19.4°C (MHWI) and 23.9°C (MHWIV) for Sines, and 18.9°C (MHWI) and 23.0°C (MHWIV) for Cabo Raso. At the end of the experiment (day 25), sea urchins were subjected to respirometry trial and subsequently sampled for morphometric analysis and gonadal tissues collection.

### Intermittent flow respirometry

Oxygen consumption rates (*M*O_2_) were measured using intermittent flow respirometry. The respirometry system consisted of individual acrylic chambers supplied with filtered and UV-sterilised seawater maintained at the corresponding experimental temperature of each treatment. Each chamber was connected to an external recirculating loop equipped with optical oxygen sensors (PyroScience, Germany), allowing continuous monitoring of dissolved oxygen concentrations. Oxygen data were recorded using AquaResp 3 software (PyroScience, Germany).

Prior to each experimental trial, optical oxygen sensors were calibrated using a FireSting optical oxygen meter (PyroScience, Germany). To account for microbial respiration, the system was run with empty chambers for approximately 15 min before each trial to quantify background oxygen consumption. Following each respirometry trial, chambers were again run empty for 15 min to reassess background respiration. Seawater was then replaced before the subsequent measurements. During measurements, chambers alternated between flush and closed phases. Flush periods lasted 5 min, during which chambers were supplied with fully oxygenated seawater, followed by 20 min closed measurement periods during which oxygen consumption was recorded. This cycle was repeated three consecutive times for each individual. Prior to each trial, chambers were inspected to ensure the absence of air bubbles and proper water circulation.

Dissolved oxygen concentrations were measured using fibre-optic oxygen sensors connected to a FireSting optical oxygen meter (PyroScience, Germany). Water recirculation and chamber flushing were controlled using external pumps connected to the respirometry system. Oxygen uptake rates were calculated from the linear decline in dissolved oxygen concentration during the closed phases and expressed as oxygen consumption rates. Given the sedentary behaviour of *P. lividus* and the difficulty in distinguishing active versus resting states, measurements were used as an estimate of overall metabolic oxygen consumption rather than resting or maximum metabolic rates.

### Morphometric measurements and gonad sampling

Sea urchins were measured for test diameter and height using a vernier caliper. Whole-body wet mass was determined using a precision balance (± 0.0001 g). Individuals were then opened orally–aborally using scissors, and gonadal tissues were carefully excised and weighed to the nearest 0.0001 g. Gonadal samples were immediately frozen on dry ice and stored at −80°C until transportation to the University of Hong Kong for molecular analyses.

### RNA extraction and sequencing

A total of 34 gonad samples from female sea urchins (4 to 6 samples per population/treatment) were used for RNA extraction and sequencing at the University of Hong Kong (Suppl. Table S2). We targeted female gonadal tissue because reproduction in *P. lividus* is particularly vulnerable to MHW-induced thermal stress (Amato et al., 2025), and female individuals have been demonstrated to be more susceptible to such events than their male counterparts (Okamoto et al., 2025). Gonadal tissue was homogenized in RLT buffer with 4-5 sterile 1.4 mm ceramic beads (Qiagen, 13113-325) using two 30s cycles at 1500 rpm with 15s intervals in a TissueLyzer (Qiagen). Total RNA was isolated using the RNeasy Mini Kit (Qiagen), following the manufacturer’s instructions, including an on-column DNA digestion step to remove potential DNA contamination. The quality of the isolated RNA was determined using a 4200 TapeStation (Agilent), and high-quality samples (RIN > 7) were sent to the Centre for PanorOmic Sciences (CPOS, HKU) for mRNA library preparation and sequencing. Stranded cDNA libraries were prepared using the KAPA mRNA HyperPrep Kit with 100ng of total RNA as input and sequenced on an Illumina NovaSeq 6000 platform to generate 150bp paired-end reads. An average of 30.9 ± 2.6 million read pairs were obtained upon sequencing across all individuals (Suppl. Table S4).

### Sequence processing and *de novo* transcriptome assembly

Raw reads quality was checked with FastQC (v0.11.8; Andrews, 2010). Illumina adapters and low quality reads were removed using Trimmomatic (v0.39; Bolger et al., 2014) with the following parameters: SLIDINGWINDOW:4:30; minimum length = 40; mismatch allowed = 2, read score paired ended = 30; read score single ended = 15; remove leading low quality or N bases below quality 8. Potential contaminant sequences from bacteria, fungi, and virus were then identified and removed with Kraken 2 (v2.1.2; Wood et al., 2019) using a confidence score of 0.5. After filtering, an average of 25.2 ± 2.4 million read pairs (Suppl. Table S4) were used for *de novo* transcriptome assembly using DRAP (v.1.92; Cabau et al., 2017) with Trinity (v2.8.4; Grabherr et al., 2011), yielding an initial assembly of 1,153,768 contigs. Transcripts containing open reading frames (ORFs) were subsequently identified using TransDecoder (v5.5.0; https://github.com/TransDecoder/TransDecoder) and only those containing ORFs ≥ 100 amino acids long were retained. These transcripts were then further filtered by BLASTP search against UniProtKB/Swiss-Prot database (-evalue 1e-5 -max_target_seqs 1), prioritizing matches first by homology and then by ORF length, resulting in 92,464 contigs. To further reduce redundancy, we used Corset (v1.09; Davidson & Oshlack, 2014) with default settings to hierarchically cluster the Transdecoder filtered contigs based on sequence similarity and expression patterns, resulting in a final transcriptome assembly consisting of 43,780 high-quality contigs. The quality and completeness of the *de novo* assembled transcriptome was evaluated with ‘TrinityStats.pl’ script from Trinity accessory scripts, rnaQUAST (v2.1.0; Bushmanova et al., 2016), ‘stats.sh script’ from BBMap (v38.87; Bushnell, 2014) and Benchmarking Universal Single-Copy Orthologs (BUSCO) analysis (v5.4.5; Simão et al., 2015) with the *metazoa_odb10* gene set (Suppl. Table S5). Annotation of the assembled transcriptome was performed using DIAMOND (Buchfink et al., 2021) in sensitive mode and UniProtKB/Swiss-Prot, UniProtKB/TrEMBL, UniRef90, and BLAST nr databases. Association of annotated genes to GO terms was performed in OmicsBox (v 3.4.5; BioBam Bioinformatics, 2019), where eggNOG mapping was also run, for a total of 27,652 successfully annotated transcripts in the final transcriptome assembly.

### Differential expression analysis

Transcript counts were obtained with Salmon (v1.10.2; Patro et al., 2017) in *quant* mode with the following specific parameters: --*gcBias*, --*libType A*. Counts were imported in R (v4.3.1; R Core Team, 2023) and used for differential gene expression analyses with DESeq2 (v1.40.2; Love et al., 2014). The presence of outliers and batch effects was evaluated by principal component analysis (PCA) and heatmaps of the sample-to-sample distances using variance stabilized transformed (VST) counts. In order to identify the genes differentially expressed because of the interaction between the treatment and the population, we ran a likelihood ratio test (LRT; design: ∼Population*Treatment, where “Treatment” has three levels: control, MHW1, MHW4). Additionally, pairwise comparisons were performed within each population between the three MHW treatment levels, and between the two populations exposed to the same heatwave treatment, using Wald tests. Significantly differentially expressed genes (DEGs) had false discovery rate (FDR) adjusted p-value < 0.05 (Benjamini & Hochberg, 1995), a mean expression of > 10 reads (baseMean) and apeglm (Zhu et al., 2019) shrunk |log2 Fold Change| > 0.3 to reduce false positives.

To identify which gene expression patterns could be correlated with either the heatwave treatment sea urchins experienced, their population of origin or any of the physiological response measured, a weighted gene co-expression network analysis (WGCNA) was performed using the WGCNA v1.73 package (Langfelder & Horvath, 2008). Physiological measurements such as test width (mm), wet weight and gonad weight (g), and oxygen consumption (mg of O_2_/g per hour) were provided as traits data. Additionally, population of origin (Cabo Raso = 1; Sines = 2), heatwave treatment (Control = 1; MHW1 = 2; MHW4 = 4) and condition, produced by the combination of the two beforementioned factors (Cabo Raso-Control = 1; Sines-Control = 2; Cabo Raso-MHW1 = 3; Sines-MHW1 = 4; Cabo Raso-MHW4 = 5; Sines-MHW4 = 6) were provided as numerically encoded trait data. The following parameters were used to build the network: power = 8 (with R^2^ > 0.90), networkType = "signed", TOMType = "signed", maxBlockSize = 9360, deepSplit = 3, reassignThreshold = 0, minModuleSize = 75, mergeCutHeight = 0.3, verbose = 3. Clusters of genes whose expression patterns were correlated with either heatwave treatment or population of origin were identified.

Enrichment analyses of differentially expressed genes identified by DESeq2, as well as gene sets belonging to significant modules detected by WGCNA, were performed with Fisher’s Exact Test (FDR < 0.05) in OmicsBox.

Finally, pathway enrichment analyses were run with Gene Set Enrichment Analysis (GSEA v4.3.2; Subramanian et al., 2005) with gene set permutation on DESeq2 normalized counts, using KEGG pathways as gene sets. Differently from classic overrepresentation analysis, GSEA analyzes all available genes without requiring an arbitrary threshold and works on the assumption that genes act in concert, where even a small change of different genes involved in the same pathway likely has stronger effects than a big fold-change modification in a single gene. Enriched gene sets were assigned based on FDR q-value < 0.05.

### SNPs calling and detection of outlier loci

Single-nucleotide polymorphisms (SNPs) calling and outlier detection was performed to determine if there were any genetic differences between the samples from the naive (Cabo Raso) and the power plant (Sines) locations. First, we mapped the high quality, decontaminated reads to the *de novo* transcriptome using bowtie2 (v2.5.4; Langmead & Salzberg, 2012) in sensitive mode not allowing discordant or individual alignments. Then, the Genome Analysis Toolkit (DePristo et al., 2011; GATK v4.5.0.0; McKenna et al., 2010) was used to flag duplicated reads in the obtained BAM files with MarkDuplicates (Picard) and split reads containing Ns in their cigar string with SplitNCigarReads with default settings. SNPs within the samples were then called using HaplotypeCaller and hard-filtered using the VariantFiltration tool (parameters: - window 35 -cluster 3 --filter-name FS -filter ‘FS>30.0’ --filter-name QD -filter ‘QD<2.0’). The resulting VCF files from all individuals were sorted and merged with bcftools (v1.13; Danecek et al., 2021). Vcftools (v0.1.16; Danecek et al., 2011) was then used to remove all indels and retain only high confidence SNPs with a Phred-scaled quality score of 30 and minor allele frequency (MAF) ≥ 0.1. Additionally, the minimum and maximum depth cut-off was set to 5 and 46, respectively, per site and per genotype to account for possible mapping/assembly errors and all sites where over 30% of the individuals are missing a genotype were removed. Discriminant Analysis of Principal Components (DAPC; Jombart et al., 2010) from the R package adegenet v2.1.10 was used to determine genetic structuring between the samples and to evaluate if variants were potentially under selection. Finally, to detect outlier loci, high quality SNPs were inspected with BayeScan (v2.1; Fischer et al., 2011) with the assumption that the individuals from the two sites belong to two different populations.

## Results

### Physiological parameters

To investigate possible effects of thermal history on the response of *Paracentrotus lividus* sea urchins to different intensity marine heatwaves, we measured morphological traits and oxygen consumption. Gonad weight to total body weight proportion was found to be significantly different between the two populations. Power plant sea urchins (Sines) in particular showed a significantly smaller gonad weight proportion than the naive population (Cabo Raso; Suppl. Figure S1; ANOVA; p-value = 0.0136). However, no significant difference was found within and between the two thermal treatments and controls.

Oxygen consumption did not differ between populations but was significantly different between treatments (Figure 2; ANOVA; p-value = 0.0011). For within population comparison, Post Hoc False Discovery Rate pairwise comparison revealed significant differences between sea urchins from the naive population kept at control and exposed to an extreme heatwave (MHW4; p-value = 0.0255), between power plant sea urchins exposed to a moderate heatwave (MHW1) versus an extreme one (MHW4; p-value = 0.0284) and between the two populations exposed to a moderate heatwave (MHW1; p-value = 0.0405). Only a non-significant mild trend for the interaction between the treatments and the two populations was found (p-value = 0.0563).

**Figure 2.**
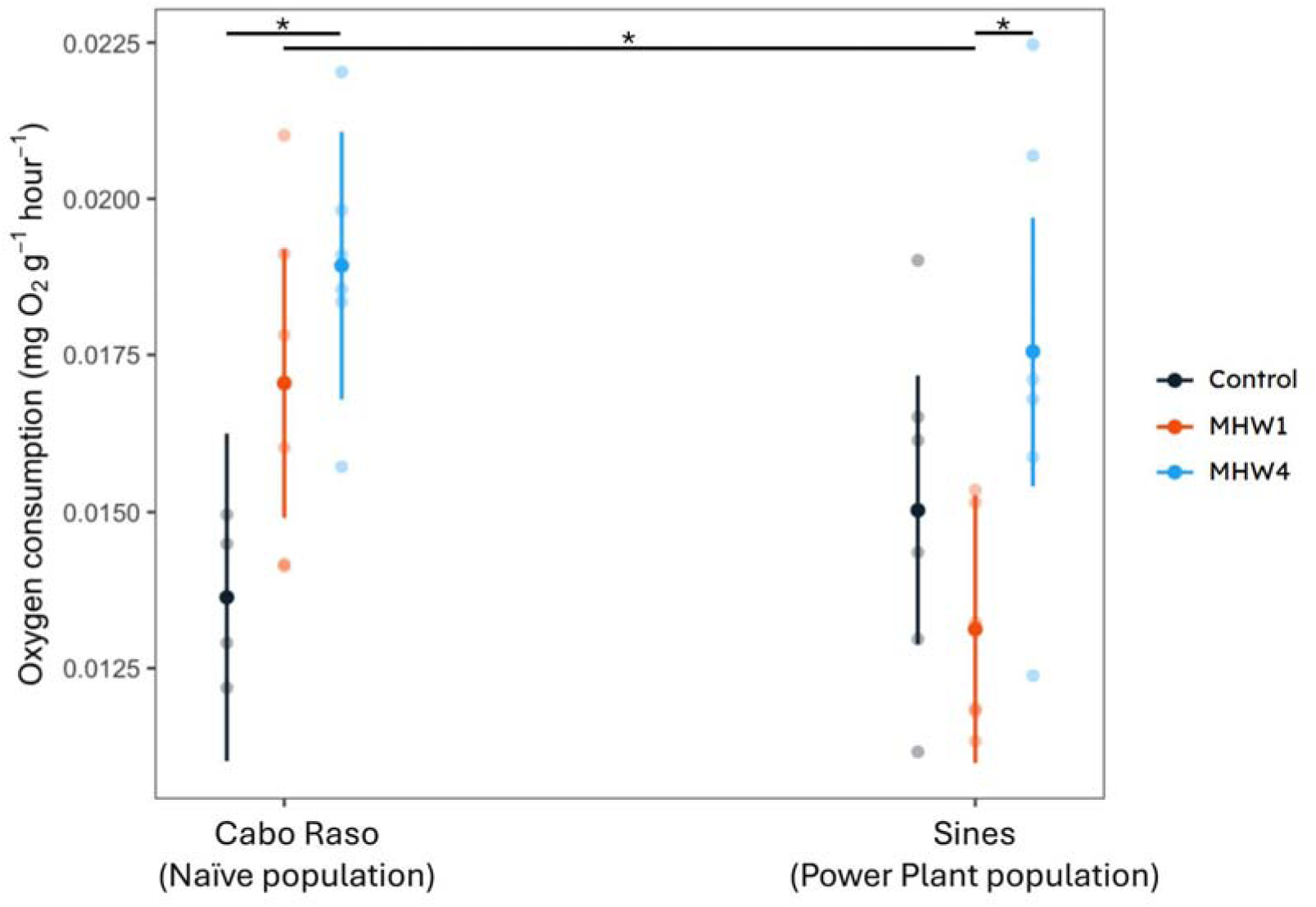
Sea urchin respiration rate differences. Exposure to heatwaves of different intensities caused significant differences in both populations in oxygen consumption. MHW1 is the moderate heatwave and MHW4 is the extreme heatwave. The asterisks show significant differences (p-value < 0.05).

### Gonad gene expression

In order to understand if the development of the power plant (Sines) population at warmer temperature has affected how they cope with warming during adulthood, we assembled *Paracentrotus lividus* female gonadal transcriptome *de novo* and compared the gene expression of these sea urchins to a naive population (Cabo Raso) developed in natural sea thermal conditions. The assembled transcriptome resulted in 43,780 contigs of N50 3,071 bp, L50 9,104 bp, with an average of 58.1% mapping rate and 97.8% of complete metazoa BUSCO genes. No genes were found to be significantly differentially expressed because of the interaction between thermal treatment and population of origin. When we ran pairwise comparisons between the two populations at different temperatures, we found the biggest difference in DEGs (123) between the two populations after the exposure to a moderate heatwave (MHW1). Conversely, when exposed to a stronger heatwave (MHW4), only 4 genes were differentially expressed between the power plant and the naive sea urchins (Fig. 3).

**Figure 3.**
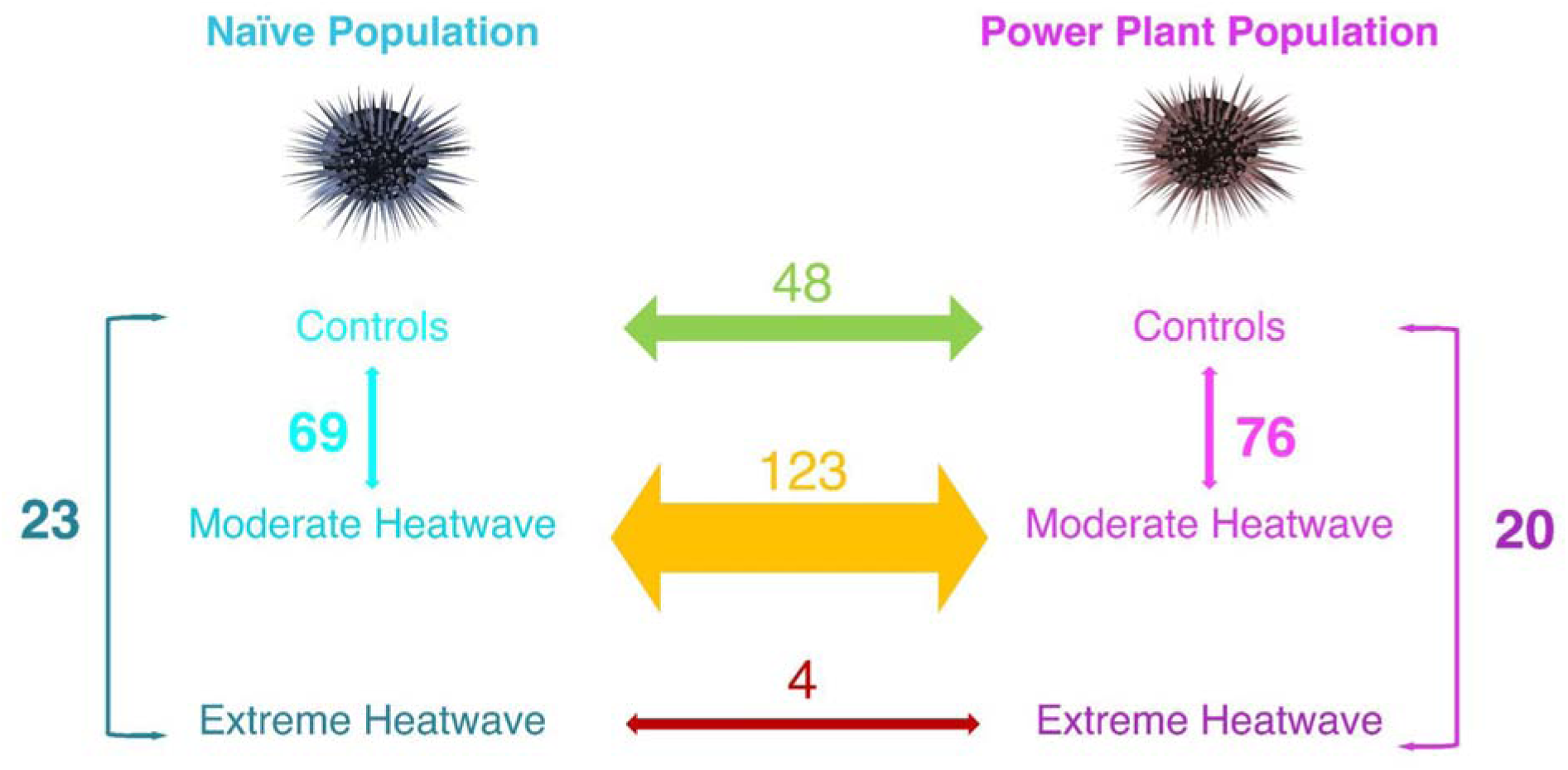
Differentially expressed genes (DEGs) between and within the two populations exposed at different heatwaves. The largest number of DEGs was found between the two populations when exposed to a MHW of moderate intensity (123 DEGs), while the smallest between the two populations when exposed to a MHW of extreme intensity (4 DEGs).

#### Population baseline differences

At control temperature, power plant (Sines) urchins showed 48 differentially expressed genes (DEGs) compared to sea urchins from the naive population (Cabo Raso; Suppl. Table S6). Most of these DEGs were upregulated in power plant individuals and included lipid transporters (e.g. apolipoproteins, vitellogenins, *hdlbp*, *fabp3*) linked to lipid transport (GO:0006869), positive regulation of lipid localization (GO:1905954) and nutrient reservoir activity (GO:0045735; Suppl. Table S7), as well as reproductive proteins (hyalin, *sfe1*). Other overexpressed genes were involved in oxidative stress response and inflammation, like glutathione S-transferases (*gstt3*, *gsta4*), solute carrier family 7 member 11 (*slc7a11*) and cytochrome P450 4F6 (*cyp4f6*), as well as immune system (e.g. *cfb, dmbt1, echinoidin*). Genes involved in innate immunity and inflammation (e.g. *grn, ltn1, nlrp1*) were also part of a module of 80 co-expressed genes (“violet” module; Suppl. Table S8; Suppl. Fig. S2) positively correlated with power plant sea urchins (p-value 0.03) and negatively correlated with gonad weight (p-value 0.04). Similarly, a module of 253 genes (“grey60” module; Suppl. Table S9) was also positively correlated with power plant urchins (p-value 0.01) and significantly enriched for a large number of different functions, such as multicellular organism development (GO:0007275), macroautophagy (GO:0016236), vesicle-mediated transport (GO:0016192) and insulin receptor signaling pathway (GO:0008286). Finally, a third module of highly correlated genes (“green” module; 1374 genes; Suppl. Table S10) was instead composed of genes expressed at lower levels in the power plant urchins compared to the naive population (p-value 0.02) and correlated with smaller size (p-value 0.04). RNA processing (GO:0006396), in particular mRNA splicing, via spliceosome (GO:0000398) and ribosome biogenesis (GO:0042254), as well as nucleosome organization (GO:0034728) and histone binding (GO:0042393) were pathways enriched in this module. Accordingly, RNA processing genes (e.g. *rcl1, slc4a1ap, u2surp*) were also differentially expressed between sea urchins from the two populations at control temperature.

#### Within and between populations response to a moderate heatwave

When exposed to a moderate heatwave (MHW1), sea urchins from one population differentially expressed 123 genes compared to the other, with only one transcript (not annotated) also DE at control temperature (Suppl. Table S11). Nevertheless, some of the differences in gonadal maturity that were present between populations at control temperature were retained during the moderate heatwave, since vitellogenin was the most strongly differentially expressed transcript. More than 80% of the DE transcripts were upregulated in individuals from the control population compared to the power plant urchins. Naive (Cabo Raso) sea urchins overexpressed stress-responsive factors including histone modifiers (e.g. *ash1l, sirt6*), DNA damage sensor *atr*, chaperone peptidylprolyl isomerase, and proteins involved in oxidative stress response (e.g. *gst1, hao1, slc7a11*) compared to power plant (Sines) urchins. They also upregulated chromatin remodelers (e.g. *chd1, paf1, rsf1*), several mobile element-related genes (reverse transcriptases, RNA-directed DNA polymerases) and cell cycle regulators (e.g. *mical3*, ribonuclease H, *tuba1c*). Moreover, an enhanced RNA splicing activity was evidenced by overexpression of splicing factors (e.g. *arglu1, cwc22, rbm19, srsf1*) and spliceosome pathway enrichment (NES −1.789, FDR q-value 0.040; Suppl. Table S12; Suppl. Fig. S3).

Within the naive population alone, the moderate heatwave caused the differential expression of 69 genes (mostly upregulated) compared to sea urchins from the same population kept at control temperature (Suppl. Table S13). These included cell cycle regulators (e.g. *mcm4, ppp4r1*), inflammation related genes (e.g. *cyp4f3, plaa, sbno1*), glutathione S-transferases (e.g. *gst1, gstt3*) and other transcripts involved in oxidative stress response (e.g. *slc7a11, uckl1*), as well as several translation factors (e.g. *eef1b, mrpl51, ptcd1, rps13, upf3b*). Pathway analysis revealed significant enrichment for spliceosome (NES 2.378, FDR q-value 0.000), proteasome (NES 2.300, FDR q-value 0.000), RNA polymerase (NES 1.882, FDR q-value 0.009), antigen processing and presentation (NES 1.745, FDR q-value 0.032), and nucleotide excision repair (NES 1.735, FDR q-value 0.031). On the other hand, the genes expressed at lower levels were enriched in pathways mostly related to immune response, like B and T cell receptor signaling pathways (NES −1.973, FDR q-value 0.032 and NES −1.863, FDR q-value 0.027, respectively), leukocyte transendothelial migration (NES −1.940, FDR q-value 0.030), and Fc gamma R-mediated phagocytosis (NES −1.870, FDR q-value 0.029), along with cell signaling pathways (MAPK: NES −1.862, FDR q-value 0.024; phosphatidylinositol: NES −1.835, FDR q-value 0.023) and adherens junction (NES 1.828, FDR q-value 0.022; Fig. 4; Suppl. Table S14).

**Figure 4.**
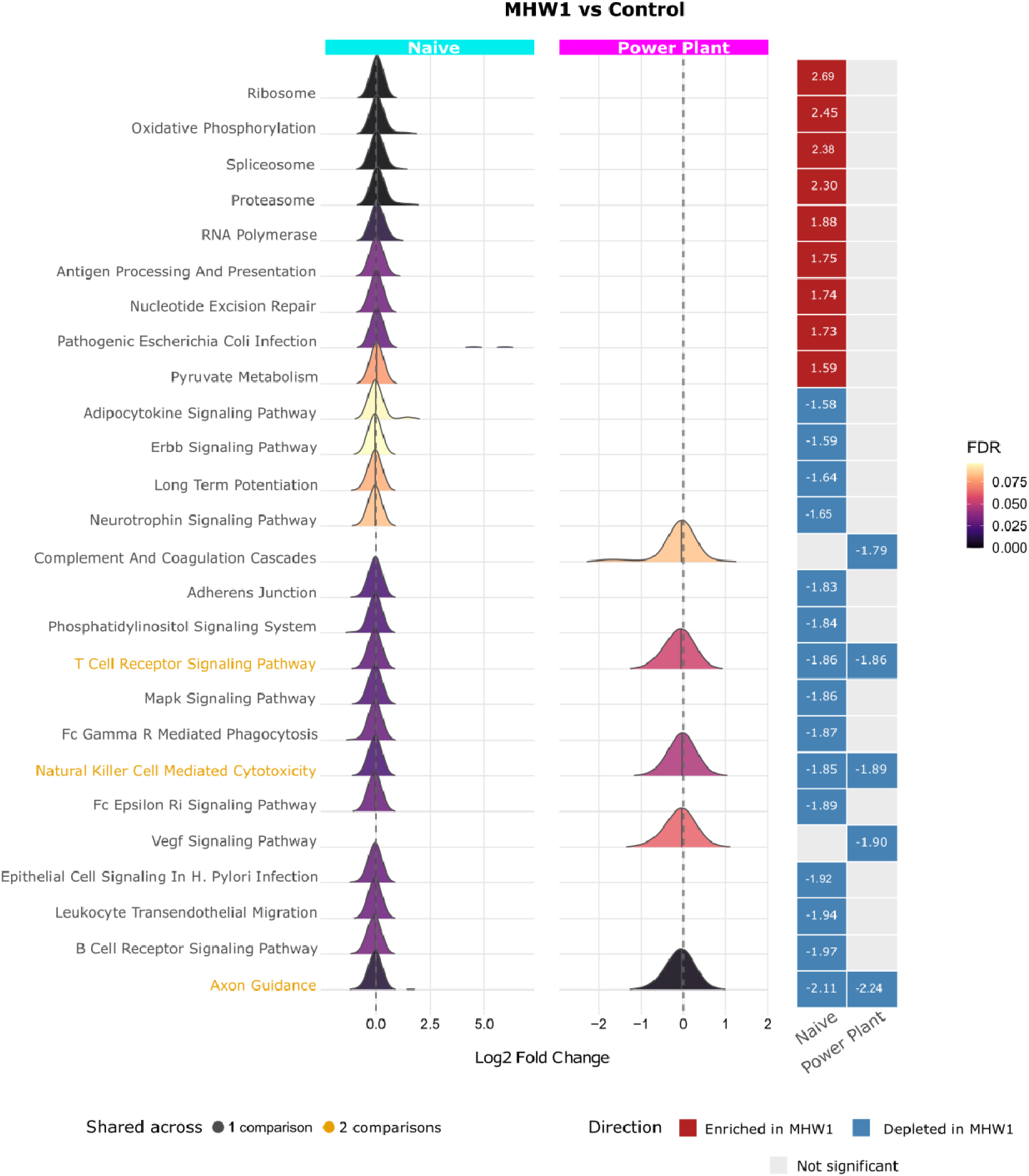
Enriched pathways in sea urchins from the two populations exposed to a moderate heatwave compared to controls. Ridge plots showing the distribution of log2 fold changes for genes belonging to significantly enriched KEGG pathways (FDR < 0.1) identified by GSEA. Each ridge represents a pathway, coloured by FDR. Pathways significant in both comparisons are highlighted in orange. The tile panel (right) summarises enrichment direction and normalised enrichment score (NES) for each comparison.

Power plant sea urchins exposed to a moderate heatwave differentially expressed 76 genes compared to power plant urchins kept at control (Suppl. Table S15). Upregulated transcripts included metabolic (e.g. *mlycd, pet100, snx10*) and gene expression (e.g. *ccdc12,* histone H4, *nr2c2ap*) regulators, as well as immune factors (e.g. *cd163*, reelin domain-containing proteins, *tnfaip8*). Conversely, downregulated genes affected proteolysis (e.g. *klhl38, uba1, xpnpep1*), translation (e.g. *celf4, ddx6, serbp1*), cell proliferation and differentiation (e.g. *fat4, fgfr, rad50*). Additionally, several genes involved in lipid transport and gonadal maturation (e.g. apolipophorins, *hdlbp*, vitellogenin) were downregulated in the exposed sea urchins compared to control samples.

#### Responses to an extreme heatwave

After an extreme heatwave (MHW4), only four transcripts were differentially expressed and upregulated in the gonads of naive (Cabo Raso) sea urchins compared to that of power plant (Sines) urchins (Suppl. Table S16). These transcripts coded for Toll like receptor 4 (*tlr4*), involved in innate immune response, fibropellin 1-like, a component of the oocyte extracellular matrix, fatty acid synthase (*fasn*), related to both lipid metabolism and inflammatory response, and an uncharacterized protein. The naive population still showed enrichment for the spliceosome pathway (NES −2.344, FDR q-value 0), similarly to after a moderate heatwave. On the other hand, power plant urchins exhibited immune and inflammatory-related pathway activation, including leukocyte transendothelial migration (NES 2.201, FDR q-value 0.001), natural killer cell mediated cytotoxicity (NES 2.096, FDR q-value 0.004), complement and coagulation cascades (NES 1.944, FDR q-value 0.011), lysosome (NES 1.884, FDR q-value 0.015), and cell adhesion molecules (CAMs; NES 1.769, FDR q-value 0.037; Suppl. Table S17).

Sea urchins from the naive population exposed to an extreme heatwave differentially expressed 23 genes compared to controls (Suppl. Table S16). Most of the transcripts were upregulated and included some genes already upregulated during a moderate heatwave, like *ltv1*, fumarate hydratase (a component of the Krebs cycle), and glutathione S-transferases, as well as new responses like ganglioside biosynthesis (e.g. alpha-N-acetylgalactosaminide alpha-2,6-sialyltransferases), heparan sulfate degradation (e.g. *naglu*), cell adhesion (e.g. *svep1*) and transcriptional regulation (e.g. BEN domain-containing protein). Again, several transcripts coding for mobile element polymerases were differentially regulated in these samples. The pathways enriched after the extreme heatwave were glycolysis/gluconeogenesis (NES 2.116, FDR q-value 0.011), fatty acid metabolism (NES 2.000, FDR q-value 0.016), antigen presentation (NES 1.933, FDR q-value 0.016), and proteasome (NES 1.950, FDR q-value 0.017; Suppl. Table S19).

Power plant sea urchins exposed to an extreme heatwave differentially expressed 20 transcripts, 18 of which upregulated, compared to same population individuals kept at control temperature (Suppl. Table S20). Among them, we found developmental regulators (e.g. choice-of-anchor I domain-containing protein, p16 protein), detoxification enzymes (e.g. *cpne7, ggt1, scarb2*), unfolded protein response activators (e.g. *xbp1*), and, similarly to the naive population, mobile element polymerases. Several immune pathways (e.g. natural killer cell mediated cytotoxicity, TLR signaling, leukocyte migration) were enriched in heatwave-exposed urchins, while RNA degradation (NES −1.891, FDR q-value 0.029) and spliceosome (NES −1.916, FDR q-value 0.042) showed negative enrichment (Suppl. Table S21).

### SNPs and outlier loci detection

A total of 7,779 high-quality SNPs were detected across all naive and power plant sea urchins. No genetic structuring was found among the analysed locations, and no outlier was detected (Fig. S4). Therefore, we found no evidence of genetic divergence between individuals collected in the two sites, with individuals from the two locations likely coming from a heterogenous pool of larvae.

## Discussion

Acclimation through phenotypic plasticity can aid organisms to cope with rapidly changing environments and may be an important mechanism for population persistence under climate change. In particular, developmental plasticity, achieved during early life stages and then maintained throughout the entire life-span, can confer advantages to marine ectotherms when facing subsequent thermal stresses (e.g. Donelson et al., 2011; Muñoz et al., 2015; Pottier et al., 2022) and is theorized to be essential for populations’ survival against unpredictable weather events linked to climate change (Burggren, 2018), such as marine heatwaves. Accordingly, here we show that a population of *P. lividus* sea urchins that developed at elevated water temperature (+2.9°C on average) exhibits acclimation signatures and responds more effectively to a simulated heatwave of moderate intensity compared to a control population. Importantly, sea urchins from the power plant site were collected nine months after the shutdown of the thermal effluent, meaning that individuals were no longer experiencing elevated temperatures at the time of sampling. Thus, the observed physiological and transcriptional differences likely reflect persistent developmental effects acquired during earlier life stages rather than short-term acclimation to ongoing thermal exposure. We may attribute these effects to epigenetic differences acquired during larvae-post settlement developmental stages rather than to genetic local adaptation, since we observed genetic homogeneity across the two populations. Physiologically, developmentally acclimated urchins displayed lower oxygen consumption under moderate heat stress compared to naive urchins. Because warming elevates oxygen demand in ectotherms, potentially leading to a mismatch between their metabolic needs and their capacity to extract and deliver adequate oxygen, a controlled respiration rate under thermal stress suggests more efficient energy use (Seebacher et al., 2015), a trait also observed in damselfish developmentally exposed to warming (Donelson et al., 2011). Accordingly, the two sea urchin populations activated different genes to cope with the moderate heatwave. However, these benefits seemed to disappear under an extreme heatwave (+5.5–6°C above control), which erased population-specific differences. This breakdown may reflect either a mismatch between expected and realized environmental conditions (“predictive adaptive response” hypothesis; Bateson et al., 2014; Hoffmann & Bridle, 2022) or that thermal limits have been exceeded, consistent with the +6°C threshold previously shown to exceed larval thermotolerance in this species (Martino et al., 2021). Since marine heatwaves of this intensity already occur in temperate regions (Wernberg et al., 2013) and are projected to increase in frequency (Huang et al., 2025), our findings suggest that extreme events may overwhelm developmental plasticity’s protective capacity, causing deleterious effects across life stages regardless of early thermal acclimation, which underlies some of the limits of adaptive developmental plasticity in a warming ocean.

While not presenting differences in terms of genetic structuring, probably due to their long pelagic larval stage (20-40 days; Fenaux et al., 1985) that allows for high gene flow over great distances (Barrier et al., 2024; Duran et al., 2004), the molecular analysis revealed that power plant urchins displayed higher constitutive expression of oxidative stress defense genes. This included glutathione S-transferases (GSTs), which conjugate reactive electrophiles to glutathione (GSH) for detoxification; solute carrier family 7 member 11 (*slc7a11*), a key transporter regulating cysteine uptake for GSH biosynthesis; and cytochrome P450 enzymes, which metabolize xenobiotics and mitigate oxidative damage. Because heat stress increases reactive oxygen species (ROS), it can overwhelm antioxidant defenses and cause oxidative stress (Slimen et al., 2014), leading to ferroptosis (Xie et al., 2021), a form of cell death by lipid peroxidation exacerbated by heat (Koppula et al., 2021; Leng et al., 2025; Stockwell, 2022). Enhancing the GSH system (supported by *slc7a11*) could help maintain redox balance by replenishing glutathione, while enzymes like GSTs and P450 can detoxify lipid peroxidation byproducts. A persistent upregulation of such detoxification pathways in thermally acclimated urchins may protect them from oxidative stress and ferroptosis (Ursini & Maiorino, 2020; Xue et al., 2025), showing their acclimation to chronic oxidative stress likely induced by the elevated temperatures near the power plant discharge. Concurrently, immune-related genes were upregulated in power plant urchins, possibly as a preemptive response to heat-induced inflammation, a pattern observed in other warming exposed organisms (Ganeshan et al., 2019; Li et al., 2025; Yang et al., 2024) and in heat-resistant *S. intermedius* sea urchins strains (Ding et al., 2017; Han et al., 2023). However, the activation of these protective pathways is energetically expensive, and might come at the cost of trade-offs. For instance, power plant urchins had smaller gonads, and immune gene expression correlated negatively with gonadal size, mirroring findings in *Xenopus tropicalis* frogs, where developmental acclimation to warming lead to shifts in energy allocation toward immune function at the expense of gonadal development and fertility (Li et al., 2025). Moreover, genes associated with gene expression, protein synthesis, and DNA organization were constitutively downregulated in power plant urchins, indicating a broad metabolic reprioritization away from reproduction toward maintaining protective and stress response functions, a potential cost of chronic thermal acclimation. Accordingly, exposure to increased temperatures negatively affects *P. lividus* energy budget and reproductive potential, with lower gonad weight relative to the somatic mass in exposed urchins (Yeruham et al., 2020). Together, these phenotypic and molecular differences highlight how developmental thermal acclimation near the power plant drove a complex suite of physiological adjustments that enhance stress resilience but may also impose significant trade-offs, potentially in reproductive investment, a life-history strategy favoring survival over fecundity in challenging environments.

The transcriptional differences between the two populations were even more evident when exposed to a moderate heatwave. Not only the number of differentially expressed genes between power plant and naive sea urchins more than doubled after the heatwave exposure, but the two populations also revealed different genes expressed in response to the heat stress, likely shaped by their differing thermal histories. Naive sea urchins, which developed under ambient temperature conditions, showed a pronounced upregulation of genes involved in stress and inflammation as well as DNA repair. Sirtuins, for example, are conserved NAD+ -dependent class III histone deacetylases crucial for managing cellular stress responses. Sirtuin 6 (*sirt6*) in particular is activated under heat stress in *Caenorhabditis elegans*, where it extends their survival through formation of stress granules (Jedrusik-Bode et al., 2013). Moreover, it acts at different levels: it reduces inflammation by blocking NF-κB target genes; controls reactive oxygen species (ROS) caused by mild stress; helps maintain cell balance by managing autophagy, which removes damaged proteins and organelles; and also repairs DNA damage by regulating chromatin proteins (Chen et al., 2025). Another key gene involved in heat stress response and upregulated in naive sea urchins during the moderate heatwave is the ataxia-telangiectasia and Rad3-related (*atr*). Under heat activation, *atr* initiates a signaling cascade to induce cell cycle arrest to prevent damaged DNA from being passed on during cell division, thereby reducing the rate of apoptosis (Furusawa et al., 2012). Accordingly, several cell cycle transcripts were also differentially regulated in naive sea urchins, suggesting tight regulation of cell division and DNA replication. Additional signals of on-going stress in naive sea urchins during the moderate heatwave were the differential regulation of genes involved in protein folding (e.g. peptidylprolyl isomerases) and the enrichment of proteasome pathway, indicating heightened protein quality control under thermal challenge. Finally, naive sea urchins also upregulated RNA splicing and other transcriptional and post-transcriptional regulatory mechanisms in response to the moderate heatwave. The activation of spliceosome has also been observed in other echinoderms, with the enrichment of spliceosome following heat stress in *S. intermedius* sea urchins (Han et al., 2023), as well as the identification of more than a thousand differential alternative splicing events in response to acute heat stress in the sea cucumber *Apostichopus japonicus* (Y. Wang et al., 2022), highlighting the significant role of alternative splicing in acute stress response in these organisms. This broad activation of cellular maintenance and stress mitigation mechanisms therefore suggests a robust acute response to heat stress, mobilizing diverse protective processes to counteract thermal damage. Notably, naive urchins downregulated immune-related pathways during the heatwave, which may reflect a reallocation of energy toward immediate stress responses at the expense of immune function.

In contrast, the transcriptional response to the moderate heatwave of sea urchins from the power plant population, which developed at chronically elevated temperatures, was characterized by differential expression of metabolic regulation and energy homeostasis genes. These developmentally thermally exposed urchins showed upregulation of key metabolic regulators including malonyl-CoA decarboxylase (*mlycd*), which promotes fatty acid oxidation through reduction of malonyl-CoA levels, and PET100 cytochrome c oxidase chaperone (*pet100*), essential for proper assembly of mitochondrial respiratory chain components. Interestingly, these urchins also upregulated sorting nexin 10 (*snx10*), a stress-responsive regulator that enhances ATP production efficiency while minimizing oxidative damage through selective control of respiratory complex turnover (Trachsel-Moncho et al., 2025). Power plant sea urchins, therefore, appear to regulate metabolic pathways to meet increased energy demands during thermal stress, mirroring patterns seen in thermally acclimated ectotherms, including heat-resistant *S. intermedius* sea urchin strains (Han et al., 2022, 2023). The moderate heatwave also triggered activation of immune homeostasis pathways in power plant urchins. For example, TNF alpha induced protein 8 (*tnfaip8*) is involved in immune responses in the sea cucumber *Apostichopus japonicus*, where it controls the metabolism of L-arginine, crucial for immune function and tissue repair, and nitric oxide content, negatively regulating inflammation (Shao et al., 2017). Similarly, the gene coding for CD163 molecule (*cd163*) also protects against oxidative stress and excessive inflammation, by acting as a phagocytic receptor (Etzerodt & Moestrup, 2013). Therefore, unlike thermally naive sea urchins that suppressed immunity during the moderate heatwave, power plant urchins maintained and even upregulated immune-related transcripts, reflecting a preconditioned state that balances cellular maintenance with metabolic and immune homeostasis. At the same time, the coordinated downregulation of proteolysis- and translation-related transcripts likely reflects another energy-regulating strategy under thermal stress. By slowing protein synthesis, cells may avoid producing proteins prone to heat-induced damage that would require degradation, thereby reducing protein misfolding or facilitating alternative folding pathways (Pechmann et al., 2013). Overall, these findings indicate that developmental exposure to elevated temperatures enables power plant urchins to mount a more efficient, metabolically adjusted stress response with sustained immunity during moderate heatwaves, while naive urchins rely on energetically costly broad protective gene activation, underscoring their vulnerability to sudden thermal stress.

The exposure to an extreme heatwave caused the homogenization of transcriptional responses of the two populations, with only four genes showing differential expression between exposed power plant and naive sea urchins, and around 20 between exposed individuals and controls within each population. Strong heatwaves with water temperature increases of +3-6°C have been found to compromise key physiological traits of sea urchins, such as gonadal development and reproduction (Amato et al., 2025), and to increase mortality (Cuthbert et al., 2021; Minuti et al., 2021). Over time, metabolic rates increase until hitting physiological thresholds that trigger metabolic collapse and death (Byrne & Hernández, 2020). For example, elevated respiration rates were documented in *Heliocidaris erythrogramma* sea urchin during heatwave conditions, followed by a decline in health index and an increase in mortality rates (Minuti et al., 2021). Here, both populations showed significantly higher oxygen consumption rates under the extreme heatwave scenario, with naive urchins also upregulating genes involved in metabolism and energy production, potentially to sustain remodeling of cell components and detoxification pathways. Indeed, a common response of to the two populations was the activation of detoxification pathways, with the upregulation of glutathione S-transferases in naive urchins and genes involved in glutathione homeostasis (e.g. *ggt1*) and scavenger receptors (e.g. *scarb2*) in power plant individuals. Additionally, both populations addressed thermal protein damage, although through distinct mechanisms. Power plant urchins activated the unfolded protein response (UPR), with the upregulation of the UPR transcriptional activator X-Box binding protein 1 (*xbp1*), also known to promote thermal injuries repair (Wang et al., 2023), while naive urchins predominantly degraded damaged and misfolded proteins via the proteasome system. The main differences between the two populations’ responses to the extreme heatwave, however, entitled modifications in cell membrane lipid composition in naive sea urchins, with the upregulation of genes coding for sialyltransferases and fatty acid synthase (*fasn*), and a sustained immune competency in power plant sea urchins, with the enrichment of multiple immune-related pathways. While the remodeling of cell surface glycoconjugates and extracellular matrix components may protect cells against heat-induced damage and preserve tissue structure during stress (Hazel, 1995), an activation of immune responses even under severe thermal stress might allow the power plant population to more efficiently counteract pathogenic bacterial infections, which become more prevalent at elevated water temperatures. Such infections have been linked to sea urchin disease outbreaks and mass mortality events under anomalously high seawater temperatures (Clemente et al., 2014; Sweet et al., 2016). Collectively, these results reveal that extreme heatwaves trigger fundamentally convergent physiological responses in both populations, which include metabolic acceleration, coordinated detoxification pathways, and protein damage mitigation systems. While variations in membrane remodeling and immune regulation persist, the overwhelming similarity in stress responses suggests these shared mechanisms constitute the core adaptive toolkit for sea urchins facing extreme heatwave events, regardless of developmental exposure to elevated water temperatures.

## Conclusion

Our findings highlight that developmental plasticity can serve as an effective means for organisms to acclimate to environmental stressors, although only up to physiological thresholds. Exposure to elevated temperatures during development can improve thermal tolerance and metabolic efficiency under moderate warming; however, its protective effect diminishes under extreme heat events. The loss of these acclimatory advantages during severe heatwaves reveals critical thresholds beyond which plasticity alone may be insufficient to ensure population persistence. This underscores the importance of considering the frequency, intensity, and duration of thermal extremes in predicting species’ responses to climate change, as well as the potential costs and trade-offs associated with chronic acclimation.

## Data Availability

The raw sequencing data and the assembled transcriptome can be found under NCBI Bioproject PRJNA1371644. Reviewer access to the BioProject record is available at: https://dataview.ncbi.nlm.nih.gov/object/PRJNA1371644?reviewer=nmc25snehmciamq6niu75mnld9. The assembled transcriptome has been deposited as a Transcriptome Shotgun Assembly project at DDBJ/EMBL/GenBank under accession GLMA00000000.

## Acknowledgements

We thank the HKU Centre for PanorOmic Sciences (CPOS).

## Authors’ contributions

T.R., J.R.P. & C.S. designed the experiment. T.R. & J.R.P. collected the animals. T.R., J.R.P and B.P.P conducted the experiment, marine heatwave exposures and respirometry assays. B.P.P. performed tissue sampling. D.R. and E.O. prepared the samples for sequencing. J.R.P analysed the physiological and morphological data. L.C.B, S.S., D.R., J.S., A.C., J.K., D.D., M.C. and J.R.P. performed the data analysis with feedback from C.S.. L.C.B. wrote the first manuscript draft, with inputs from all the authors. T.R. and C.S. secured the funding. All authors read, provided comments and gave final approval for publication.

## Supplementary Figures

**Supplementary Figure S1.**
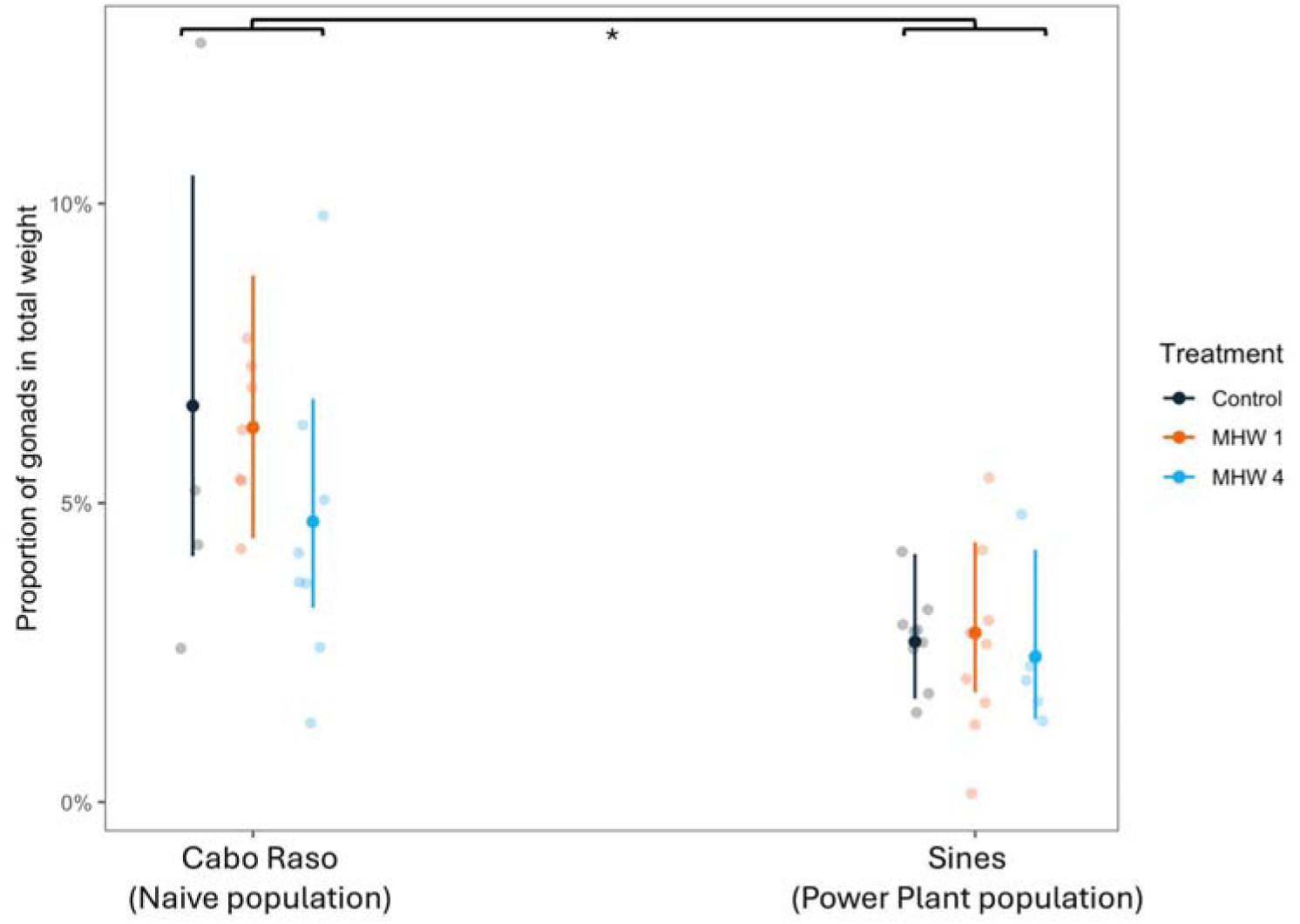
Differences in the proportion of gonads in total weight between the populations. Naive (Cabo Raso) sea urchins had a significantly bigger gonad to total weight proportion than power plant sea urchins that developed at higher temperatures (Sines). MHW1 stands for moderate heatwave, MHW4 for extreme heatwave. The asterisk shows the significant difference (p-value < 0.05).

**Supplementary Figure S2.**
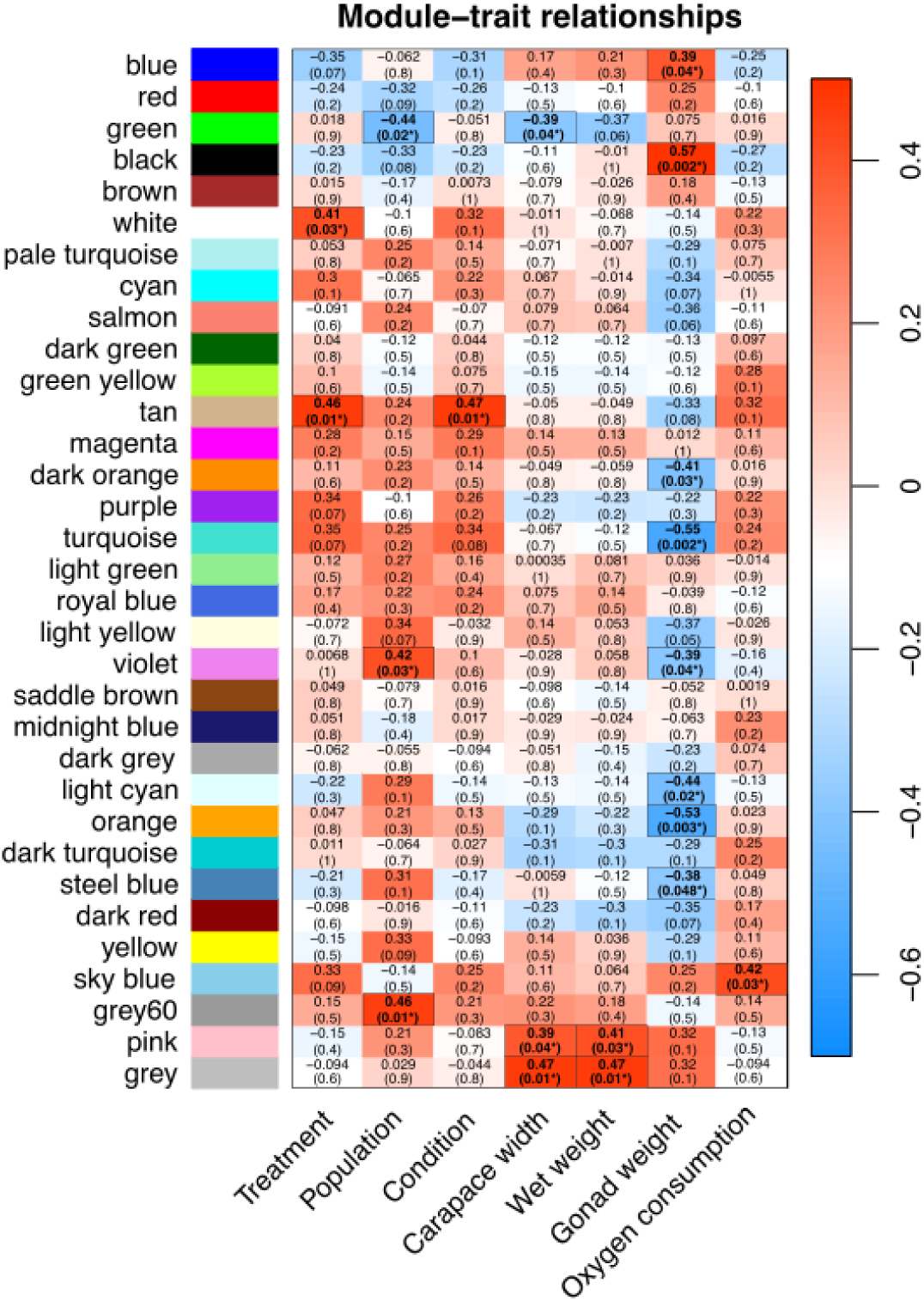
Heatmap of module-trait correlation analysis between detected gene network modules (denoted by an arbitrary color name), warming treatments, population of origin and physiological parameters. In each cell is reported the correlation value for the corresponding module eigengenes and each treatment and the p-value (in brackets).

**Supplementary Figure S3.**
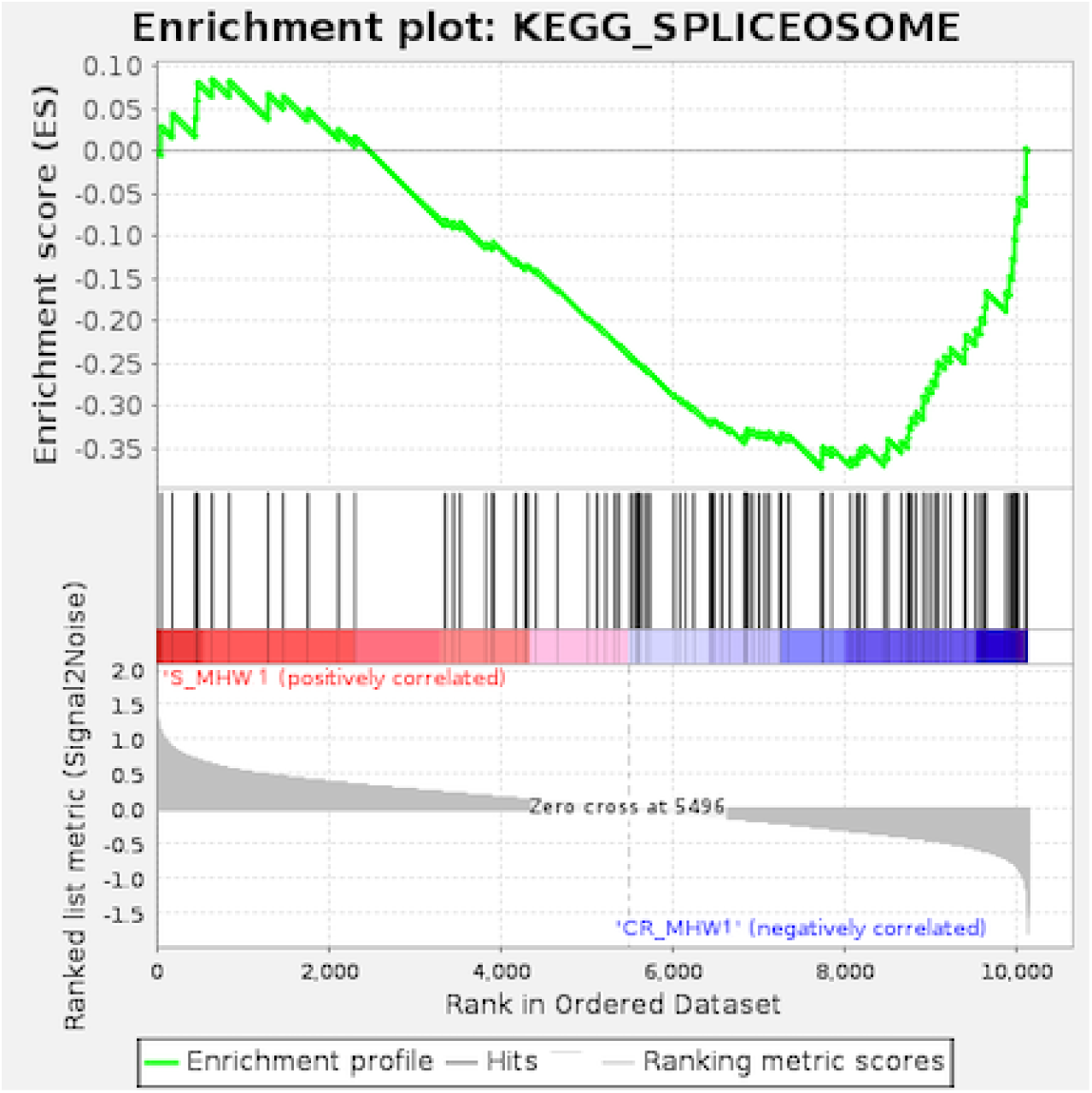
Enrichment plot for “Spliceosome” KEGG gene set in sea urchins from the power plant population (Sines - S) compared to the naive one (Cabo Raso - CR) under a moderate heatwave (MHW1).

**Supplementary Figure S4.**
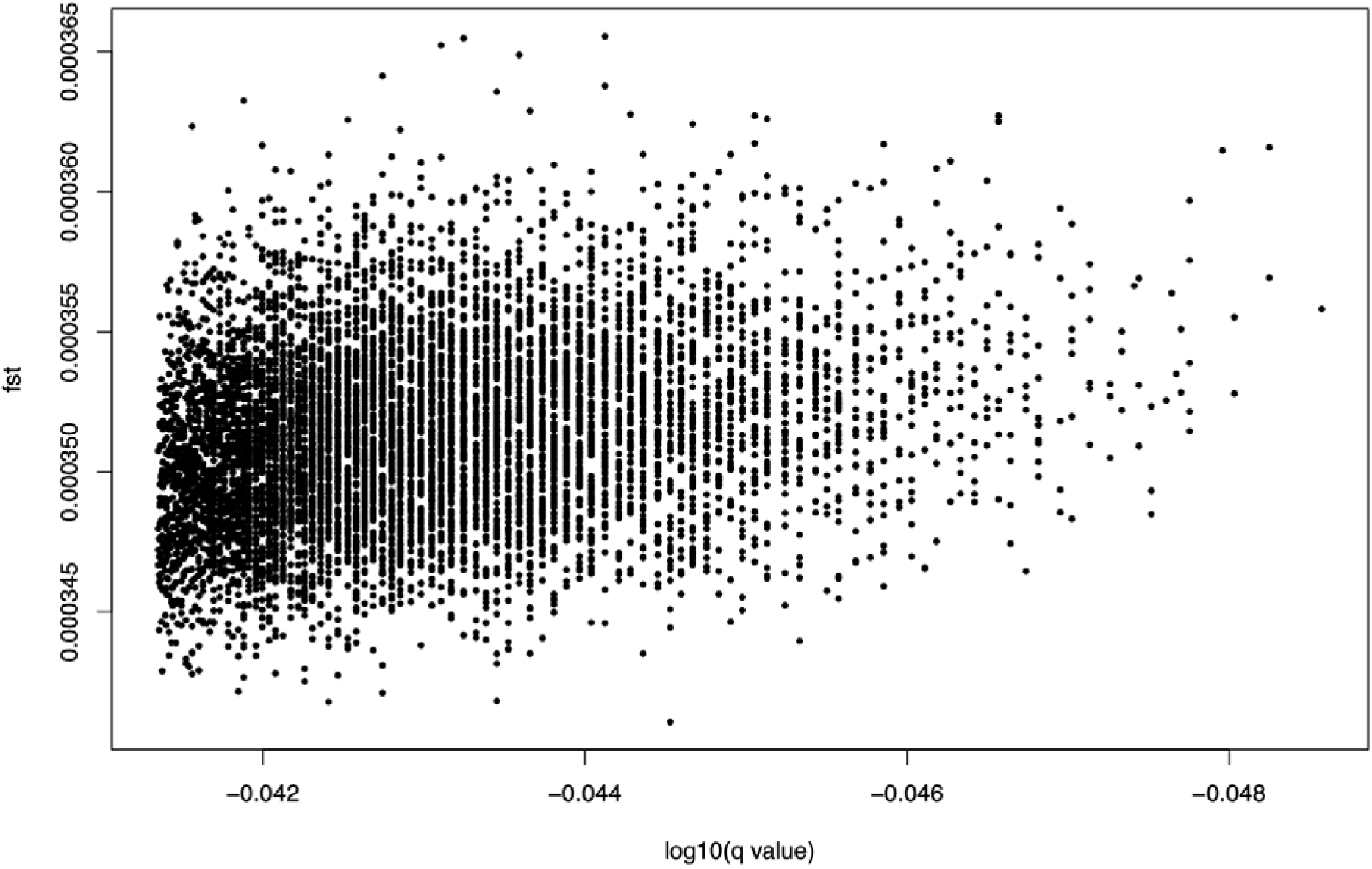
Results of Bayescan analysis for 7,779 SNPs detected between individuals from the two locations. The graph represents the F_ST_ values against the corresponding log10(FDR corrected p-value(q value)) for each loci. No outlier loci identified.

